# Rising novelty and homogenization of breeding bird communities in the U.S.

**DOI:** 10.1101/2022.09.27.509749

**Authors:** C. E. Latimer, R. A. Graves, A. M. Pidgeon, J. M. Gorzo, M. Henschell, P. R. Schilke, M. L. Hobi, A. Olah, C.M. Kennedy, B. Zuckerberg, V. C. Radeloff

## Abstract

**Aim:** Human modification has profound effects on the diversity of ecological communities. Yet, surprisingly little is known about how abiotic novelty due to human modification relates to biological novelty as measured by shifts in species composition from historical baselines. Using space-for-time substitution, we ask a) whether high human modification results in biotic homogenization or heterogenization across different spatial scales; b) if high modification results in the formation of novel, “no-analog” communities; and c) whether changes in bird community composition varies in response to proxies of historical land-use and duration-of-exposure to anthropogenic disturbances.

**Location:** Conterminous United States.

**Time Period:** 2012 – 2016.

**Major taxa studied:** Passeriformes.

**Methods:** We analyzed continent-wide avian biodiversity data from an online checklist program, eBird, to examine how shifts in breeding bird species composition have been impacted by human modification at regional and continental scales and tested four hypotheses related to how abiotic novelty resulting from human modification generates biological novelty.

**Results:** At regional scales, bird communities in highly human-modified areas exhibited similar levels of β-diversity as those in the least modified areas. However, at the continental scale, spatial turnover in community composition was lower in human-modified areas, suggesting that anthropogenic disturbance has a strong homogenizing effect on bird communities at that scale. Lastly, human modification contributed more to community composition in regions where natural disturbance was infrequent and Euro-American settlement occurred later, consistent with the hypothesis that exposure to historical disturbances can shape how contemporary bird communities respond to human modification.

**Main conclusions:** The observed patterns of increased biotic novelty and homogenization in regions with less frequent disturbances and a longer history of human modification suggests that future extensive human modification could result in further homogenization of bird communities, particularly in the western U.S. We argue that current human-modified environments hold great potential for biodiversity conservation.

## Introduction

“*Then the coal company came with the world’s largest shovel; and they tortured the timber and stripped all the land. Well, they dug for their coal till the land was forsaken; then they wrote it all down as the progress of man*.” ∼ John Prine

Human activities have caused extensive modification of habitats globally (Kennedy *et al*., 2019), which often outpace or constrain species’ abilities to appropriately respond to environmental changes (Robertson *et al*., 2013) and can lead to differences in species assemblages among sites (β-diversity; Whittaker, 1972; Karp *et al*., 2012). However, empirical evidence of the impact of human disturbance on the composition of species assemblages and, specifically, on local and regional beta-diversity is mixed. In some cases, human disturbance causes the convergence or homogenization of biotic communities (lower β-diversity; Mckinney & Lockwood, 1999; Olden & Rooney, 2006), but in others, it causes divergence or heterogenization (increased β-diversity; Laurance *et al*., 2007; Tscharntke *et al*., 2012; Socolar *et al*., 2016). These conflicting outcomes may be due to the spatial variation of anthropogenic disturbances, and the degree to which they produce abiotic conditions that are outside the historical range of environmental heterogeneity (i.e., abiotic novelty) (de Castro Solar *et al*., 2015; Radeloff *et al*., 2015b; Lindenmayer *et al*., 2019), in addition to the spatial scale of investigation (Karp *et al*., 2012; McGill *et al*., 2015). For instance, if human landscape modifications generate similar environmental conditions over vast areas, that tends to favor generalist species with strong dispersal abilities, and results in taxonomic homogenization (Olden & Poff, 2003). However, if human disturbances create additional heterogeneity at finer spatial scales, that can lead to taxonomic diversification (Tscharntke *et al*., 2012; Stein *et al*., 2014; Socolar *et al*., 2016; Chase *et al*., 2019). In addition to changing environmental heterogeneity, human disturbance produces novel abiotic conditions, and our question here is whether that leads to the creation of novel biotic communities (Radeloff *et al*., 2015a; Soininen *et al*., 2018; Heger *et al*., 2019). Improved understanding of the scale-dependent relationships between community dissimilarity and landscape modification can help assess the degree to which human disturbance results in biotic novelty and elucidate the mechanisms underpinning changes in biotic communities, thereby informing biodiversity conservation strategies in the Anthropocene.

Beta-diversity depends on different mechanisms at local and regional scales (Barton *et al*., 2013). At local scales, species richness can increase rapidly due to variation in species responses to anthropogenic disturbances (Barton *et al*., 2013). Consequently, ecological filters that cause differences in species richness between sites likely cause increases in *β*-diversity among sites within regions (Svenning *et al*., 2011). At regional scales, however, *β*-diversity is more likely constrained by historical dynamics (e.g., glaciation, biogeography) causing differences in ecological sorting across environmental gradients (Baselga, 2010a; Svenning *et al*., 2011; Betts *et al*., 2019).

Deconstructing *β*-diversity into its two constituent components (i.e., species replacement and species loss) helps to understand the mechanisms causing scale-dependent community responses to landscape modifications (Baselga, 2010; Svenning *et al*., 2011; Karp *et al*., 2012). Species replacement, or turnover, occurs in response to human modification, for example, when specialist species are replaced by generalists in the most intensively-altered areas without a concomitant change in species richness (Baselga, 2010; Socolar *et al*., 2016). Nestedness describes the degree of loss or extirpation of species that cannot tolerate human modification. Thus, nestedness generates patterns in *β*-diversity independent of species turnover (Baselga, 2010). The two components of *β*-diversity reflect separate processes, generated by different mechanisms (Baselga, 2010), and with different conservation implications (Socolar *et al*., 2016).

We deconstructed *β*-diversity into turnover and nestedness to identify mechanisms driving North American bird *β*-diversity at regional and continental scales. We focused on birds because they are widespread, ecologically diverse and have experienced consistent and widespread declines over the past half century (Rosenberg *et al*., 2019). We compared bird communities along human modification gradients, and formulated and evaluated four complementary hypotheses about the role of human modification as a driver of structure and composition of bird communities at regional and continental scales (Figure 1):

**Figure 1:**
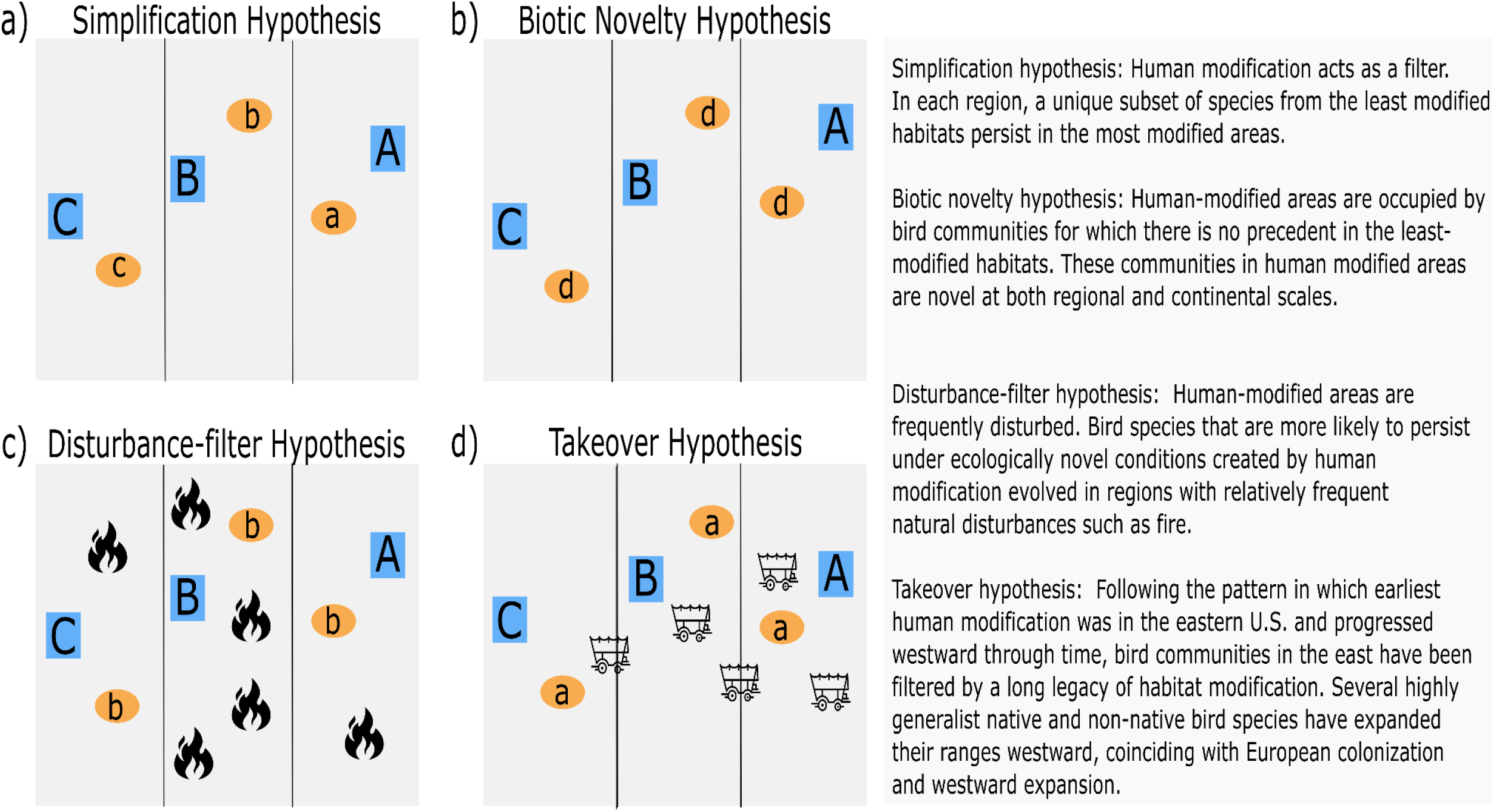
Conceptual diagram representing the four hypotheses tested in this study (panels a – d). Blue squares with upper-case lettering represent hypothetical bird communities within low human modified areas unique to each region. Orange colored circles with lower-case lettering represent hypothetical bird communities within human modified areas. Lower-case lettering denotes expectations about which region human modified bird communities most closely resemble those of more natural communities. A lower-case “d” indicates a completely novel hypothetical community.

1. *Simplification hypothesis*: This hypothesis predicts that human-modification acts as a filter, selecting a subset of the species occurring in the least-modified habitats that can persist in anthropogenic habitats (Patterson, 1987; Blair, 1996; Batary et al., 2017). Thus, simplification predicts that species richness is lower in highly modified habitats and that *β*-diversity arises primarily through nestedness of species assemblages along gradients of human modification. At broad scales, if the species persisting in the most modified areas differ among regions, then simplification will lead to heterogenization of modified bird communities among regions (e.g., *β*-diversity in highly modified habitats equal to or higher than in low-modified habitats across regions but not within regions). However, if those species persisting in anthropogenic habitats are common among regions, then simplification will lead to homogenization among regions (Figure 1a).
2. *Biotic novelty hypothesis:* This hypothesis predicts that highly human-modified areas that are abiotically novel are now occupied by bird communities for which there is no precedent in the least-modified habitats (Barnagaud et al., 2014; Lindenmayer et al., 2019; Sales et al., 2020). In other words, bird communities in human-dominated landscapes are novel both within- and among regions (Figure 1b). Both processes of turnover and nestedness could contribute to high *β*-diversity between the least- and most modified areas within a given region. However, when comparing among regions, this hypothesis predicts that bird assemblages in highly modified habitats would exhibit lower spatial turnover and higher nestedness than communities in the least modified habitats.
3. *Disturbance filter hypothesis*: This hypothesis predicts that that species’ responses to human modification depends on the frequency of exposure to natural disturbances in which the species evolved (Henle *et al*., 2004; Martin & Fahrig, 2016; Betts *et al*., 2019). For example, species that evolved in landscapes with less frequent disturbance are expected to be more at risk to anthropogenic disturbance (Martin & Fahrig, 2016). In contrast, species that evolved in frequently-disturbed habitats, such as fire-prone oak-savanna woodlands and grasslands that are often ephemeral and patchily distributed, are likely less sensitive and may even benefit from human modification (Boulinier *et al*., 2001; Schlossberg *et al*., 2011). Therefore, bird communities that historically relied upon frequent natural disturbances, such as fire, to create and maintain open habitats may be better-able to persist in human-modified areas where sparse vegetation cover resembling that of open temperate biomes is common (Figure 1c). Accordingly, if bird communities inhabiting regions characterized by more frequent natural disturbances are in fact predisposed to be synanthropic (Johnston 2001), then we expect that less of the compositional turnover would be explained by human modification in regions with more frequent natural disturbances prior to major human modification.
4. *Takeover hypothesis*: This hypothesis predicts that bird communities in the regions where widespread human modification occurs first “take over” communities in regions that are modified later, because the species in the early-modified regions had more time to adapt. As European settlement of the U.S. proceeded from east to west roughly from 1607 to1887, avian communities in the east have been subjected to a nearly 300-year longer legacy of human modification including pronounced periods of deforestation and reforestation (Whitney, 1994). Initial deforestation likely would have filtered out species that were unable to persist in human-modified environments, leading to initially higher levels of compositional turnover along gradients of human modification (subtractive heterogenization; Socolar *et al*. 2016). However, over time, generalists and invaders would likely dominate many communities (Olden & Poff, 2003), and with reforestation, some species might colonize human-altered landscapes (Corlett, 2016). In addition, several exotic species, including rock pigeons (*Columba livia*), house sparrows (*Passer domesticus*), European starlings (*Sturnus vulgaris*) and Monk parakeets (*Myiopsitta monachus*) were introduced in the Eastern U.S. and have since expanded their ranges westward (Burgio et al., 2020; Cabe 2020; Lowther & Cink 2020; Lowther & Johnston, 2020). Similarly, some native species, including the American robin (*Turdus migratorius*), chipping sparrow (*Spizella passerina*), and to a lesser extent, Northern cardinal (*Cardinalis cardinalis*), have progressively expanded their breeding ranges westward, partly owing to the establishment and intensification of farmlands and urban areas (Halkin et al., 2021; Middleton, 2020; Vanderhoff et al., 2020). Combined, we expected that these processes would have generated patterns of lower turnover along natural-modified gradients in the eastern U.S. relative to the western U.S. (Figure 1d). Therefore, under this hypothesis, we expect to see an increasing contribution of human modification and geographic distance to assemblage turnover in regions with shorter historical legacies of time since European colonization and westward expansion.

## Methods

### Bird occurrence data

We compiled information on bird occurrences from the eBird database (Version 1.11; Sullivan *et al*., 2014) for 2012 - 2016. We selected checklists for the conterminous U.S. and completed under the travelling, stationary, and area sampling protocols. For traveling, stationary, and area protocols, we limited analyses to checklists with durations of 3 hours or less and that were ≤ 3 km in length (for traveling), or ≤ 1km^2^ (for area) (La Sorte *et al*., 2014; Johnston *et al*., 2019). We only considered complete checklists in which all species seen or heard were recorded during daylight hours (05:00h – 20:00h), and that were submitted during the height of avian breeding season (June – August). We included all species from checklists (resident and migratory, and native and non-native), except species that are nocturnal or associated primarily with freshwater or marine environments. We excluded nocturnal and freshwater/marine species based on taxonomy (i.e., family) and expert review. This resulted in 1,122,084 eBird checklists available for analyses from 298,844 unique locations.

### Environmental predictors

We assigned each checklist to one of the 30 Bird Conservation Regions (BCR) represented in the conterminous U.S. (NABCI; Figure 2a). Each BCR contains internally similar bird communities, habitats and resource management issues (Bird Studies Canada and NABCI, 2014). Because our primary focus was to quantify the degree of dissimilarity in community composition resulting from human modification, we used the human modification model (Theobald, 2013) and calculated the average human modification score within a 3-km radius buffer of each eBird checklist location. Human modification scores represent the cumulative degree of land modified within 30-m pixels, ranging from 0.0 (low) to 1.0 (high), and are based on the spatial extent and intensity of urbanization, agriculture, energy production, nighttime lights and roads (Theobald 2013). Because bird richness and composition may also be affected by habitat heterogeneity within a region (Fahrig *et al*., 2011; Farwell *et al*., 2020), we calculated Shannon’s index of diversity (SHDI) of different land cover types within a 3-km radius buffer using the 2016 National Land Cover Dataset (Homer *et al*., 2019). We also extracted elevation from the National Elevation Dataset (Gesch *et al*., 2002), and maximum temperature in the hottest month (Fick & Hijmans, 2017) within the same buffer. Maximum temperature during the hottest month captures temperature extremes during the breeding season which can strongly affect occurrences of many North American bird species (Cohen *et al*., 2020). Because many checklist locations were within close proximity to one another, and to avoid potential errors associated with the location information reported by eBird participants (Zuckerberg et al., 2016), prior to analyses we tallied species from all checklists within a 3-km pixel (U.S. Albers equal-area projection) and converted them to presences (1) or absences (0), referred to hereafter as species assemblage pixels. Aggregating checklists within each pixel, as we have done here, minimizes the possibility of overinflating estimates of β-diversity due to, for example, variation among observers in their ability to detect rare species, or sampling in different habitats and years (šizling *et al*., 2011; Keil *et al*., 2012). For similar reasons, we opted to use incidence-based measures of community dissimilarity (see below), and not abundances, because the raw eBird count data exhibited high positive skew that likely would have artificially inflated estimates of turnover and nestedness. This resulted in 106,565 assemblage pixels for analyses (see Table S1, Appendix S1 for break-down by region).

**Figure 2:**
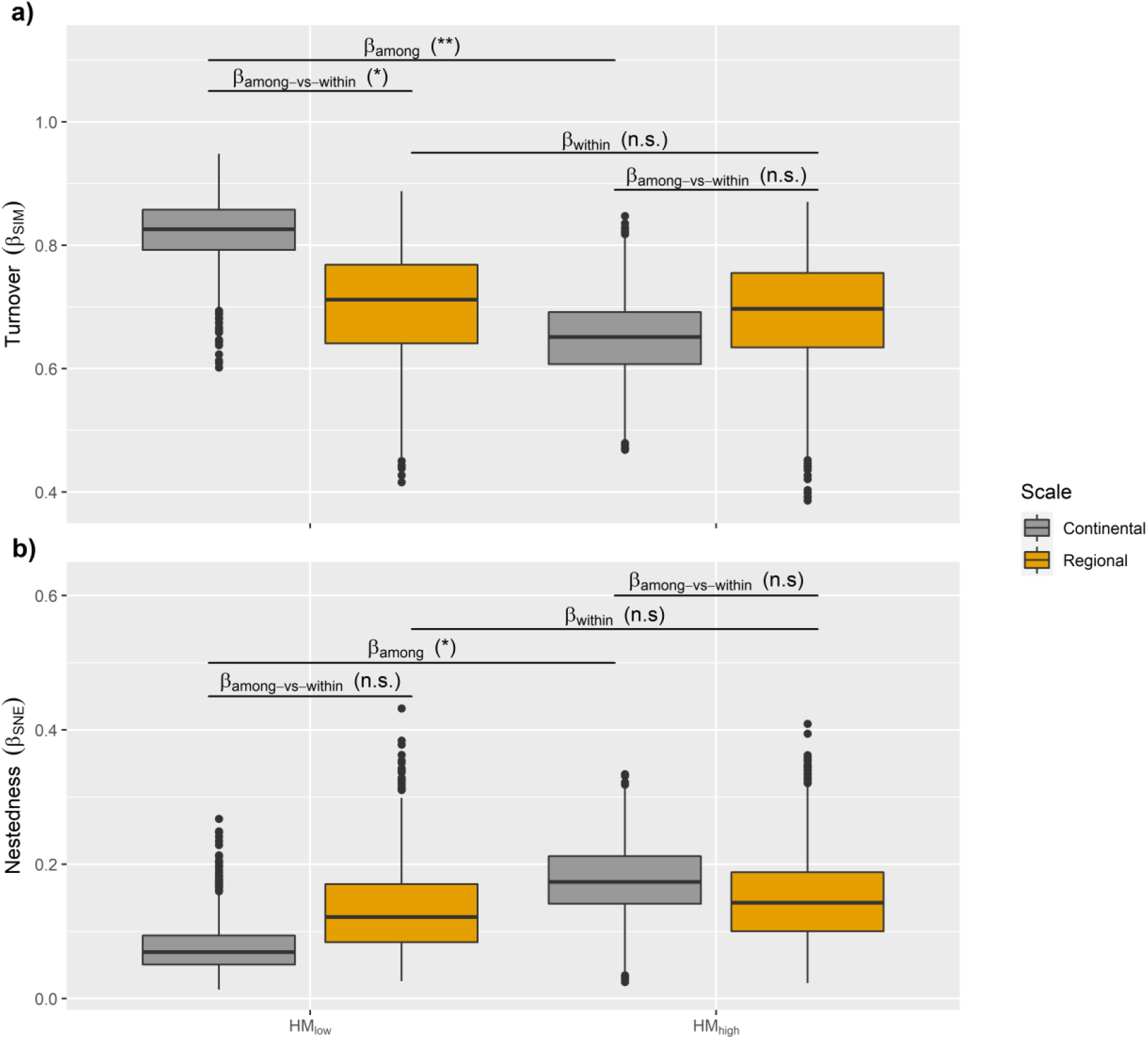
Comparisons of compositional turnover (a) and nestedness (b), within (β_within_) and between areas (β_among_) of high and low human modification at regional (within Bird Conservation Regions) and continental (among Bird Conservation Regions) scales.

To test the simplification and biotic novelty hypotheses, we chose the lower 5^th^ and upper 95^th^ percentiles of the human modification score as thresholds to represent the least (HM_low_, referred to as “Natural”) and most (HM_high_, referred to as “Modified”) human-modified pixels that were sampled by eBird participants, respectively. We calculated those percentiles separately for each BCR. We estimated the sampling coverage completeness using the total number of checklists within each human modification category in each BCR (Chao & Jost, 2012). In all cases, sampling completeness was > 95% (Fig. S1 in Appendix S1). Therefore, we could use raw species richness values to compare bird communities in landscapes with high human modification (HM_high_) to those in landscapes with low human modification (HM_low_). We note that we focused on the extreme ends of the human modification gradient because the ‘simplification’ and ‘biotic novelty’ hypotheses are primarily focused on comparisons between the least and most modified regions. In contrast, our tests of the ‘disturbance filter’ and ‘takeover’ hypotheses focus on community changes along gradients of human modification (see below).

### Statistical Analyses

#### Decomposition of β-diversity

We calculated total dissimilarity of bird communities (i.e., species assemblage pixels, as explained above) both *within* and *among* BCRs. We decomposed *β*-diversity into turnover (species replacement) and nestedness (species gain/loss), using the betapart package in R (Baselga & Orme, 2012). We calculated the total multiple-site Sorensen dissimilarity (*β*_SOR_) for each of the HM_high_ and HM_low_ groupings. Spatial turnover, independent of species richness, was the multiple-site Simpson dissimilarity (*β*_SIM_) (Baselga, 2010). Nestedness was the difference between Sorensen and Simpson indices (*β*_NES_ = *β*_SOR_ − *β*_SIM_).

Multiple-site dissimilarities are preferred over taking the average of pairwise dissimilarities because they account for species co-occurrences at more than two sites (Diserud & Odegaard, 2007; Baselga, 2010). However, multiple-site measures can be sensitive to sample size variation if regions, such as our BCRs, have different sizes (Baselga, 2010). We therefore calculated *β*-values for HM_high_ and HM_low_ groupings using a resampling procedure to differentiate between regional and continental scales. To calculate regional scale *β*-diversity (*β*_within-regions_), we randomly sampled 10 species assemblage pixels for each human modification category within each BCR, and did so 1000 times. Similarly, to calculate continental scale *β*-diversity (*β*_among-regions_), we randomly sampled 10 species assemblage pixels for each human modification category, 1000 times across all BCRs. For both within- and among-region comparisons, we empirically compared the resulting distributions of multiple-site dissimilarities for each human modification category by assessing how many times HM_high_ was higher or lower than HM_low,_ divided by the total number of samples (Baselga, 2010; Baselga *et al*., 2015).

### Tests of Hypotheses

#### Patterns of community changes in response to human modification (simplification hypothesis and biotic novelty hypothesis)

##### 1) Simplification hypothesis

We determined whether bird communities in landscapes with high human modification represent a simplified subset of those in the least modified areas using two approaches. First, we parameterized linear mixed-effects models to assess whether species richness was lower in HM_high_ than in HM_low_ regions of a BCR by including human modification as a categorical fixed effect and BCR as a random effect and a Gaussian error distribution. Second, we examined the proportional contribution of nestedness of *β*-diversity (*β*_SNE_) to overall multiple-site 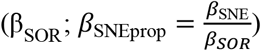 simplification hypothesis, the relative importance of local extinctions is greater than that of species replacement in determining differences in *β*-diversity between HM_high_ and HM_low_, then within a given BCR, *β*_SNEprop_ should be higher in HM_high_ regions. To test this, we used the resampling test described above and compared the distributions of *β*_SNEprop_ between HM_high_ and HM_low_ areas within each BCR.

##### 2) Biotic novelty hypothesis

To determine whether highly modified landscapes are occupied by bird communities for which there is no precedent in the least modified landscapes (i.e., novel communities), we identified a critical threshold of dissimilarity that discriminates between low and high human modification groupings based on a receiver operating characteristic (ROC) curve analysis (Gavin *et al*., 2003; Stralberg *et al*., 2009). This analysis compares the dissimilarity between each species assemblage pixel and its *k* closest neighbors within groups (analogues) to the *k* closest dissimilarities between groups (non-analogues). If the dissimilarities of samples within each of the HM_high_ and HM_low_ groupings is small compared to the dissimilarities among groupings, then the coefficient identified by the ROC analysis distinguishes between HM_high_ and HM_low_ assemblages while minimizing the false positive error rate (Gavin *et al*., 2003). We determined the optimal dissimilarity threshold value (*z*) separately for each BCR and counted the number of HM_low_ analogues based on the number of pixel assemblages for which pairwise *β*_sor_ dissimilarities were less than *z*. If there were no analogues between HM_low_ and HM_high_ groupings, then we considered bird communities present within highly modified habitats as completely novel relative to bird communities present within the least human-modified areas within a given BCR.

#### Biogeographic drivers of community novelty (disturbance-filter hypothesis and takeover hypothesis)

To evaluate our two hypotheses about the degree to which bird communities in human-modified landscapes converged with bird communities native to disturbance-prone landscapes (i.e., disturbance filter hypothesis), or in line with the historical legacy of human modification (i.e., takeover hypothesis), we calculated pairwise dissimilarities between all species assemblage pixels within each BCR. We then used generalized dissimilarity modeling (GDM) to examine how patterns of community turnover (*β*_sim_) varied along gradients of human modification, and to determine the relative contribution of various environmental and geographic drivers to avian community novelty (Ferrier & Guisan, 2006). GDM is an extension of matrix regression, suitable for modeling nonlinear patterns in compositional dissimilarity between pairs of locations as a function of environmental and geographic dissimilarity (Fitzpatrick *et al*., 2013). Through the use of flexible I-splines, GDM overcomes two major problems in community modeling: non-linear relationships between β-diversity and environmental dissimilarity, and uneven rates of changes in β-diversity along environmental gradients (Ferrier & Guisan, 2006; Fitzpatrick *et al*., 2013). I-splines are partial regression fits that serve as an indication of the importance of each variable in determining patterns of β-diversity. As such, the maximum height of each I-spline represents the total amount of compositional turnover or nestedness associated with each variable, holding all other variables constant (Fitzpatrick *et al*., 2013).

We fit separate GDMs for each BCR using a geographic distance matrix, calculated as the Euclidean distance between the centroid of each species assemblage pixel to the centroid of all other pixels, and untransformed vectors of human modification, Shannon’s diversity index, elevation and the maximum temperature during the hottest month using the R package *gdm* (Fitzpatrick *et al*., 2020). We used the default setting of three I-spline basis functions per environmental predictor and tested variable significance through Monte Carlo sampling and stepwise backward elimination (Fitzpatrick *et al*., 2013). We performed 500 permutations per step until only significant variables (α = 0.05) remained. Lastly, we determined variable importance by summing the I-spline coefficients for each environmental predictor and assessed model fit using the percent deviance explained by each GDM (Fitzpatrick *et al*., 2013). We calculated the summed I-spline coefficients from the final models to examine the overall contribution of human modification to *β*_sim_ for each BCR, after accounting for potential differences caused by variation in temperature, elevation, habitat diversity and geographic distances among pixel assemblages. We then used the summed I-spline coefficients for human modification to evaluate the two remaining hypotheses.

##### 3) Disturbance-filter hypothesis

If bird communities native to disturbance-prone ecosystems are less sensitive to anthropogenic disturbances, then we expected a lower overall contribution of human modification to turnover in regions with higher rates of disturbances (Figure 1c). As a proxy for disturbance frequency, we used LANDFIRE’s Mean Fire Return Interval (MFRI) product (LANDFIRE Biophysical Settings, 2010; V1.2.0). MFRI quantifies the average period between fires under the presumed historical fire regime, whereby shorter return intervals indicate more frequent disturbances. We calculated the median MFRI value within each BCR using GoogleEarth Engine, to obtain an overall estimate of disturbance frequency for a given BCR. Based on this dataset, large regions in both the northeast and western U.S. have longer MFRI, indicating less frequent disturbances. Under the disturbance-filter hypothesis, we expected a positive relationship between the contribution of human modification to turnover and the median MFRI value within a given BCR, indicating a greater influence of human-driven disturbance on bird community shifts in areas with lower rates of natural disturbance. We evaluated this relationship by calculating Pearson’s correlation coefficients (*r*) between the summed I-spline coefficients for human modification and the median MFRI value within each BCR.

##### 4) Takeover hypothesis

If the order in which regions were settled by Euro-Americans was important in structuring present-day bird communities, then we expected an increasing contribution of human modification to turnover in regions with a longer legacy of industrial human land-use (Figure 1d). To test this hypothesis, we used a data set on U.S. tribal land cessions, compiled by Charles C. Royce and published as the second part of the two-part Eighteenth Annual Report of the Bureau of American Ethnology to the Secretary of the Smithsonian Institution, 1896 – 1807 (USDAFS, 2018). This geospatial data set provides the name of each Native American tribe affected, date, and location of the treaty, law or executive order governing the land cessions to the U.S. government, and is based on historical records dating back to 1776. Between 1776 and 1887, the U.S. seized more than 600 million hectares of land belonging to Native Americans. We calculated Pearson’s correlation coefficients (*r*) between the summed I-spline coefficients for human modification and the median year of Native American Land cessions to the U.S. government for each BCR. A positive relationship between the summed I-spline coefficients for human modification and the median year in which Tribal lands were ceded to the U.S. government for a given BCR would suggest a greater overall contribution of human modification to *β*-diversity in regions that were colonized later. Finally, if a longer history of exposure to anthropogenic land-use change has led to greater homogenization of bird communities in the east, we expected to see a greater overall contribution of geographic distance to *β*-values in western regions, suggesting that more-distant communities in the east are more similar to each other (homogenized) than more-distant communities in the west.

## Results

### Patterns of community changes in response to human modification Simplification hypothesis

We did not find support for the simplification hypothesis. At a regional scale, species richness was lower, on average, in the most human modified landscapes (β = -0.068, S.E. = 0.01, z = -3.51, *P* = < 0.005). However, neither turnover (β_SIM_) nor nestedness (*β*_SNE_ or *β*_SNEprop_) differed between highly-modified (HM_high_) and low-modified (HM_low_) areas within a given BCR, although *β*_SIM_ was the dominant driver of bird community differences (Table 1; Figure 2a, b; *β*_within_).

**Table 1.** Overall multi-site beta diversity (β_SOR_), multi-site turnover (β_SIM_), multi-site nestedness (β_SNE_), and the proportion of β_SOR_ due to β_SNE_ (β_SNEprop_) averaged over all Bird Conservation Regions. Standard errors are in parentheses. Values are for all 3 × 3 km species assemblage pixels within a BCR (Table S1, Appendix1), the most natural pixels only (5^th^ percentile of human modification), and the most modified pixels only (95^th^ percentile of human modification).

| | $\beta_{\text{SOR}}$ | $\beta_{\text{SIM}}$ | $\beta_{\text{SNE}}$ | $\beta_{\text{SNEprop}}$ |
| --- | --- | --- | --- | --- |
| All | 0.941 (0.006) | 0.879 (0.011) | 0.061 (0.005) | 0.068 (0.007) |
| Natural | 0.938 (0.010) | 0.883 (0.016) | 0.055 (0.007) | 0.062 (0.009) |
| Modified | 0.944 (0.007) | 0.876 (0.015) | 0.067 (0.009) | 0.074 (0.010) |

At a continental scale, β_SIM_ was also the predominant driver of *β*_among-region_ (Table 1; Figure 2a, b), but turnover was higher in the least-modified areas across the conterminous U.S. (Figure 2a, *β*_among_). Moreover, in the least-modified areas among BCRs, spatial variation in species turnover (β_SIM_) was higher than that within BCRs (Figure 2a, *β*_among-vs-within_). In contrast, the spatial variation in assemblage composition due to multi-site nestedness (*β*_SNE_) was greater in highly-modified landscapes (Fig 2b, *β*_among_). Collectively, these results suggest that at broader spatial scales (i.e., among BCRs), highly-modified habitats exhibited stronger patterns of homogenization, whereas at finer spatial scales (within BCRs), bird communities in highly modified landscapes maintained similar levels of spatial variation in turnover and nestedness to those in least-modified landscapes. Therefore, bird communities in highly-modified regions do not simply resemble subsets of bird communities in least-modified regions.

### Biotic novelty hypothesis

We also did not find support for the biotic novelty hypothesis that predicts that human modification leads to novel (or “no analog”) communities across all regions within the U.S. because analog communities existed in both the least- and most-modified landscapes in all BCRs. However, the proportion of analog pixel-assemblages within BCRs ranged between 5% (high novelty) and 60% (low novelty), suggesting large variation in the degree of biotic novelty among BCRs (Figure 3a, Table S1, Appendix S1). This implies that other factors, such as biogeographic history and historical exposure to disturbances, are likely important drivers of the novelty of bird communities.

**Figure 3:**
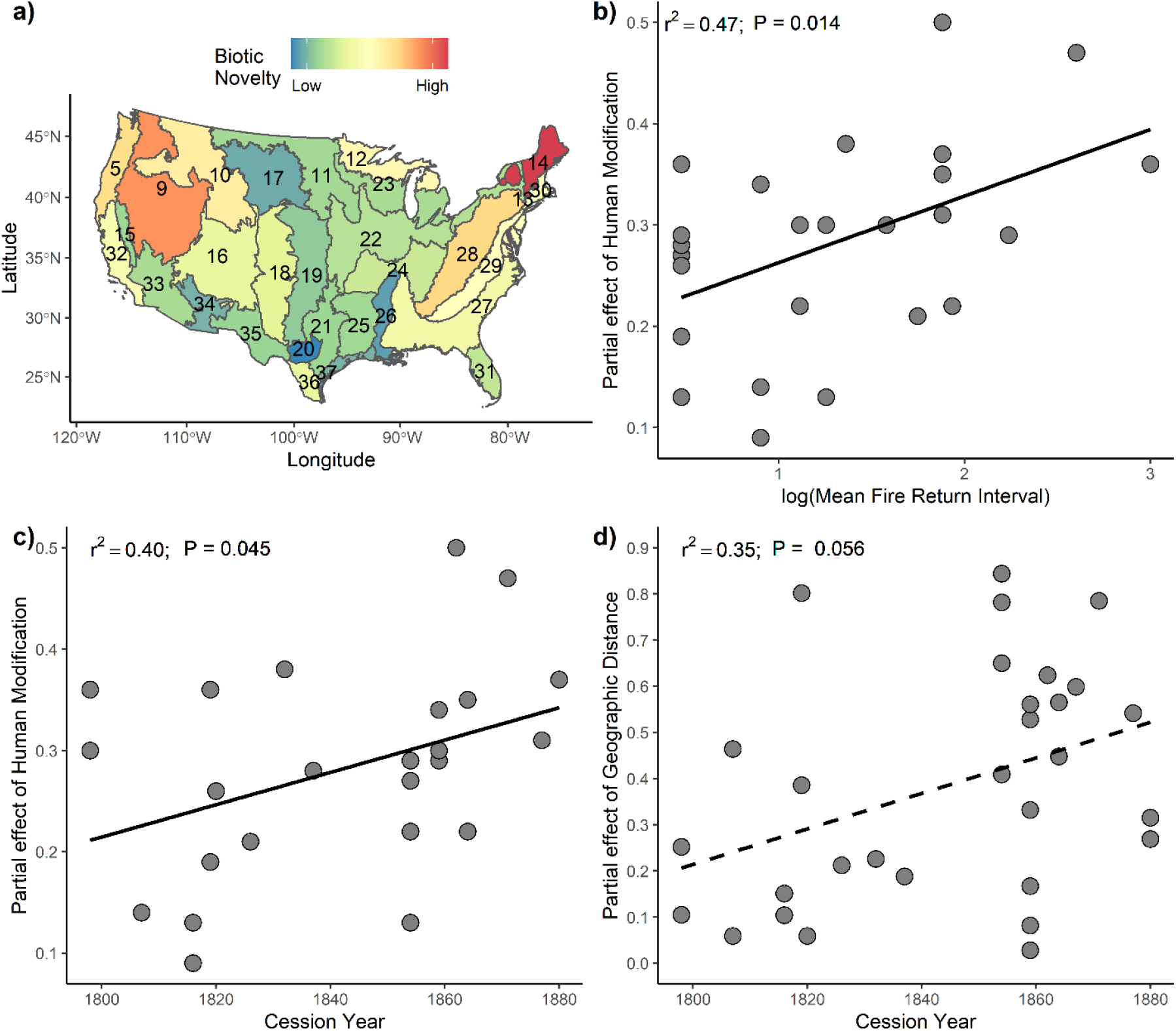
a) The proportion of analogues identified between HM_high_ and HM_low_ assemblage pixels within each Bird Conservation Region (BCR) outlined in dark gray and labeled with their numeric identifier. Proportions were inversed and rescaled between 0 (low) and 1(high) for graphical representation (see Table S1, Appendix S1 for additional details); b) the partial effect of human modification as a function of the Mean Fire Return Interval within each BCR, extracted from generalized dissimilarity models (GDM’s); c) the partial effect of human modification as a function of the median date that Native American lands were ceded to the U.S. government within each BCR; and d) the partial effect of geographic distance between pixel assemblages versus the longitudinal midpoint of each BCR. Solid regression line represents significant relationship at the α = 0.05 level; dashed regression line indicates marginal significance at the α = 0.09 level.

### Biogeographic drivers of community novelty

#### Disturbance-filter hypothesis

We did find support for the disturbance-filter hypothesis that bird communities residing in frequently disturbed habitats may be less sensitive to human modification, in that our models showed a significant positive relationship between the contribution of human modification to turnover and the mean fire return interval of each BCR (*r*^*2*^ = 0.47; *P*< 0.05; Figure 3b; Figure S2). This was true even after accounting for potentially confounding variables such as geographic distance, habitat heterogeneity and elevation. Deviance explained by these other environmental variables and geographical distances varied strongly among the 30 BCRs and ranged from 4 – 36 % (Table 2). Unsurprisingly, geographic distance was a significant predictor of pair-wise turnover in all BCRs, and human modification was retained in 26 (87%) of the final GDMs after applying the stepwise backward elimination procedure (Table 2). In addition, patterns of species turnover varied by environmental gradients and geographical distance. Elevation and geographic distance were the most important (as determined by the summed I-spline coefficients from the GDM models) predictors of turnover in 12 (40%) and 11 (36%) of the BCRs respectively. Therefore, while human modification was a significant predictor of avian compositional dissimilarity in most BCRs, it was the predominant driver in only three (Eastern Tallgrass Prairie [BCR 22], Prairie Hardwood Transition [BCR 23] and Mississippi Alluvial Valley [BCR 26]; Table 2). Shannon’s diversity, a measure of habitat heterogeneity, was kept in only 6 (20%) of the final GDMs and was the most important predictor of turnover in just two BCRs (Central Hardwoods [BCR 24] and Tamaulipan Brushlands [BCR 36]; Table 2).

**Table 2.** Relative importance of predictor variables for avian beta turnover within each Bird Conservation Region (BCR). Relative importance was determined by summing the I-spline coefficients from the final generalized dissimilarity model (GDM) after applying a stepwise backwards elimination procedure. The most important predictors for each BCR are shown in bold, and predictors found to not be significant are indicated by dashes. Deviance explained by the final model (DevExp) is also listed. Abbreviations are: Geog = Geographical distance; Elev = Elevation; Tmax = maximum temperature during the hottest month; SHDI = Shannon’s diversity index; HM = Human modification score.

| BCR | Geog | Elev | T<sub>max</sub> | SHDI | HM | DevExp |
| --- | --- | --- | --- | --- | --- | --- |
| 5 | **0.596** | 0.553 | 0.477 | - | 0.496 | 7.996 |
| 9 | 0.615 | - | 0.701 | 0.373 | 0.224 | 5.806 |
| 10 | **0.783** | - | 0.435 | 0.287 | 0.347 | 5.804 |
| 11 | **0.467** | - | - | - | 0.344 | 4.237 |
| 12 | **0.726** | - | - | - | 0.290 | 4.101 |
| 13 | 0.069 | **0.358** | - | - | - | 22.090 |
| 14 | 0.192 | **0.382** | 0.347 | - | 0.364 | 6.396 |
| 15 | 0.110 | **1.281** | 0.880 | - | 0.296 | 16.516 |
| 16 | 0.200 | **1.287** | - | 0.377 | 0.311 | 16.599 |
| 17 | 0.294 | **1.386** | - | - | - | 12.200 |
| 18 | **0.897** | 0.603 | - | - | 0.128 | 7.177 |
| 19 | **1.150** | - | - | - | 0.218 | 19.898 |
| 20 | **0.284** | - | - | - | - | 23.444 |
| 21 | **0.287** | 0.434 | - | - | 0.273 | 9.132 |
| 22 | 0.125 | 0.034 | - | - | **0.281** | 27.527 |
| 23 | 0.282 | - | 0.038 | - | **0.378** | 22.890 |
| 24 | 0.084 | 0.280 | - | **0.364** | 0.256 | 7.116 |
| 25 | 0.799 | - | **0.977** | - | 0.286 | 5.988 |
| 26 | 0.100 | - | - | - | **0.209** | 18.682 |
| 27 | 0.089 | **0.777** | - | - | 0.088 | 6.174 |
| 28 | **0.491** | 0.239 | 0.349 | - | 0.141 | 4.555 |
| 29 | 0.092 | **0.626** | - | - | 0.134 | 33.143 |
| 30 | **0.325** | - | - | 0.140 | 0.300 | 4.927 |
| 31 | **0.371** | - | - | - | 0.188 | 36.342 |
| 32 | 0.164 | **0.975** | 0.262 | - | 0.305 | 11.614 |
| 33 | 1.193 | **1.407** | - | - | 0.467 | 15.566 |
| 34 | 0.349 | **1.323** | - | - | - | 16.424 |
| 35 | 0.337 | **1.026** | 0.583 | - | 0.374 | 19.229 |
| 36 | 0.183 | 0.321 | - | **0.503** | 0.301 | 21.274 |
| 37 | 0.652 | **0.709** | - | - | 0.356 | 20.975 |

#### Takeover hypothesis

We found support for the takeover hypothesis in that the contribution of human modification to turnover increased significantly in regions with shorter legacies of intensive human modification. Furthermore, the influence of human modification, in the form of biotic homogenization, was greater in regions with a longer history of intensive human land use (*r*^*2*^ = 0.43; *P* < 0.05; Figure 3c; Figure S3). Lastly, we also found a marginal increase in the effect of geographic distance on β-diversity in regions with sorter legacies of intensive land-use, suggesting that bird communities have become increasingly homogenized through time, because the overall contribution of geographic distance to turnover was lower in regions with longer legacies of intensive modification (*r*^*2*^ = 0.31; *P* < 0.1; Figure 3d; Figure S4). As the general pattern of land cessions occurred from east to west across the U.S., this suggests that distant communities in the eastern U.S. are more similar to each other than are distant communities in the western U.S.

## Discussion

There are many open questions about how human-driven disturbance relates to biological novelty as measured by shifts in species composition that are outside of historical baselines (Belote *et al*., 2020). We decomposed beta-diversity into turnover and nestedness components and found that human modification of habitats has strong impacts on patterns of avian assemblage composition across regional and continental scales.

### Effect of human modification on avian assemblage composition

One of the consequences of human-driven biotic homogenization is that bird communities residing in more human-modified areas are more similar (have lower β-diversity) than in less modified areas (Mckinney & Lockwood, 1999). Yet, this pattern is less clear when comparing across spatial scales (Sax & Gaines, 2003). Here, we found evidence that within regions, bird assemblages in highly human-modified areas had similar levels of β-diversity, including β_SOR,_ β_SIM_, *β*_SNE_ and *β*_SNEprop_, as those in more natural areas. On the other hand, across the continent, we found strong evidence that assemblages in highly modified areas were more like one another, having less spatial variation in turnover and greater contributions of nestedness, resulting in overall lower levels of β-diversity than assemblages in least-modified areas.

Together, our results suggest that within BCRs, human modification does not simply filter a subset of species from more natural habitats, but rather, selects for a new set of species that can persist in anthropogenic habitats. There are several potential explanations for our findings. First, at narrower regional scales, human modification may be driven by one or a few land uses that create additional spatial heterogeneity, promoting greater taxonomic diversity within highly-modified areas (Fahrig *et al*., 2011; Stein *et al*., 2014). However, at broader spatial scales, more land uses are sampled, and fewer new species are likely encountered among sampling units within highly modified habitats due to deterministic variation in how species respond to human modification (Socolar *et al*., 2016). Alternatively, within regions, highly modified areas could maintain similarly high levels of β-diversity as low-modified areas, through random community assembly processes (neutral effects) or species spillover across habitats. This is opposite of what would be expected through environmental filtering, whereby greater environmental heterogeneity at broader spatial scales would lead to increases in resource complementarity and a concomitant increase in β-diversity in low-modified environments (Karp *et al*., 2012; Tscharntke *et al*., 2012). Given that we consistently detected similar patterns across both components of β-diversity that control for variation in α-diversity (Baselga, 2010), the former explanation is probably more likely. Regardless, our results suggest that within regions of the U.S., human-modified lands may hold substantial potential for conservation and the preservation of vital ecosystem functions, particularly through enhancing local diversification and maintaining matrix habitats to promote landscape-scale heterogeneity for biodiversity (Fahrig *et al*., 2011; Kremen & Merenlender, 2018). Across the continent, however, conservation efforts in human-modified landscapes may be better suited to focusing on the most diverse regions.

### Effect of human modification on biological novelty

Altered environmental conditions resulting from human modifications will differentially affect species within original communities of natural systems, favoring species that tolerate or thrive under human-dominated conditions (i.e., synanthropes) (McKinney, 2006; Sofaer *et al*., 2020; Williams *et al*., 2020). Thus, if human activities shift the composition of ecological communities towards more synanthropic species, highly modified landscapes may consist of entirely unique combinations of species, representing novel or “no-analog” communities (Williams & Jackson, 2007). We found that within regions, highly modified habitats supported communities with similar compositions as those in the least modified habitats, even though similarity differed among regions. In other words, within each region, at least one analog community could be found between those residing in the least and most modified habitats. However, regions varied substantially in the proportion of analog communities in the least and most modified habitats, ranging from low (5%) to high (60%), indicating high and low degrees of novelty, respectively. Several mechanisms could be responsible for the observed differences in regional novelty, including natural disturbance regimes such as fire (Drapeau et al., 2016) or climate dynamics (Stralberg et al., 2009), and past trajectories of land-use change (Lindenmayer et al., 2019): all of which could contribute to how contemporary bird communities respond to human modification through biotic filtering or evolutionary adaptation (Betts et al., 2019). Yet, without more temporally-resolved spatial data sets, these mechanisms are difficult to tease apart.

### Historical legacies explain how communities respond to human modification

Historical references are often used to evaluate relative changes in species persistence or ecosystem conditions and to set targets for ecological restoration (Hobbs *et al*., 2009; Lindenmayer *et al*., 2019). Yet, empirical baselines are notoriously difficult to quantify, making space-for-time substitutions to examine departures from reference conditions necessary (Newbold *et al*., 2015). We assumed that bird communities residing in the least modified habitats were representative of historical communities of natural systems, to examine how community composition shifts along gradients of anthropogenic modification. Because regions differed in their degree of biological novelty, we tested two non-mutually exclusive hypotheses (disturbance-filter and takeover hypotheses) about how patterns of community shifts and biotic novelty along human modification gradients may relate to historical land-use and prior exposure to natural disturbances. First, the disturbance-filter hypothesis, which tests if species’ evolutionary history shapes their capacity to respond to novel environmental stressors, and predicts that species that evolved in naturally-fragmented, frequent-disturbance environments are more likely to persist in human-dominated environments (Balmford, 1996; Betts *et al*., 2019).

Thus, we expected the contribution of human modification to compositional turnover would be lower in regions with more frequent natural disturbances. Using Mean Fire Return Interval as our proxy to classify regions based on their historical exposure to natural disturbances, we found a strong positive relationship between the relative contribution of human modification driving community compositional shifts and regions with high amounts of frequently-disturbed habitat. Specifically, bird communities residing in regions characterized by high-frequency disturbances experienced less turnover along human modification gradients than those within less frequently disturbed regions. This aligns with recent findings that global extinction risk from habitat fragmentation was up to three times higher in regions with low rates of historical disturbances (Betts *et al*., 2019), and supports the notion that bird communities inhabiting regions characterized by more frequent disturbances may be less sensitive to novel disturbances, including human modification (Johnston, 2001). While several mechanisms could be invoked to explain this pattern, it is likely that disturbance-sensitive species would have either gone locally extinct or adapted to frequent disturbances long before our study took place, thus shaping the response of present-day bird communities to contemporary human modification (Drapeau et al., 2016; Betts et al., 2019)

Second, the takeover hypothesis relates variation in biological novelty among regions to human modification according to the history of human modification so that regions that have been modified longer are expected to have higher levels of biotic homogenization (Balmford, 1996). This means that areas with longer legacies of industrialized and higher-intensity human activities should support greater proportions of synanthropic species, and the contribution of human modification to community compositional shifts should be lower in regions with longer histories of intensive human occupancy (Balmford, 1996). We used a geospatial database providing the date and location of the treaty, law or executive order governing tribal land cessions to the U.S. government as a proxy for length of time since Euro-American settlement and westward expansion. Over roughly three centuries (from 1607 to 1887), the U.S. seized over 600 million hectares of Indigenous lands, which generally occurred from east to west (Figure S3), and coincided with the conversion of large swaths of forests, savannas, and prairies to croplands, followed by rapid human population growth (Turner *et al*., 1980; Waisanen and Bliss, 2002). Matching our predictions, the contribution of human modification was significantly lower in regions of the United States that had a longer legacy of intensive human land use, after accounting for variation in geographic distance and other environmental factors (e.g., climate, topography and land cover diversity). Moreover, the contribution of geographic distance to dissimilarity in community composition was lower in regions with a longer legacy of intensive land-use, typically occurring in the eastern U.S. Together, these results suggest that bird communities in the eastern U.S. are more homogenized than those in the west. To our knowledge, this is the first example of how the regional order of Euro-American colonization shaped the responses of contemporary bird communities to human modification, and begs the question as to what would present-day bird communities look like had the western coast been colonized first? While several mechanisms have generated contemporary avian assemblages, the westward expansion of native and non-native generalist species breeding ranges following the establishment and intensification of urban areas and farmsteads (Cabe, 2020; Halkin & Linville, 2020; Middleton, 2020; Vanderhoff et al., 2020); and behavioral or evolutionary adaptations to human-modification (Sol et al., 2013; McCabe et al., 2018) appears to be important among them. Indeed, Sofaer *et al*., (2020) report similar findings, whereby greater proportional abundances of human-associated passerines occur within the eastern U.S. If true, this suggests that time lags between habitat loss or degradation and species extinctions may be masking the full extent to which human modification will impact bird communities in the west, and more community changes are to be expected. In turn, this could threaten key ecosystem services that birds supply, like pest-control and pollination (e.g., biodiversity-based ecosystem service debts; Isbell *et al*., 2015; Ziter *et al*., 2017).

### Limitations

Several potential limitations should be considered when interpreting our results. First, we used a cumulative measure of human modification that integrates five types of anthropogenic stressors: human settlement, agriculture, roads, mining and electrical infrastructure (Theobald, 2013). As species likely respond in different ways to each of these stressors, it could be that no-analog communities are more likely to exist within areas subject to one type of human land uses (e.g., urban areas) than another (e.g., cropland or pasture) (Newbold *et al*., 2016). Teasing apart these relationships would require further analysis of the dominant drivers of anthropogenic impacts, which was beyond the scope of our study. Second, we analyzed species presences and absences, not abundances, which would underestimate the patterns of novelty related to species’ abundances or their functional and phylogenetic diversity (Graham *et al*., 2017; Sofaer *et al*., 2020; Sol *et al*., 2020). While it is possible to obtain abundance data from eBird, abundances for many species exhibit a high right skew, suggesting that abundance-based diversity indices would have resulted in higher dissimilarity among pixel assemblages and potentially even stronger effects than we observed with incidence-based measures of dissimilarity. Third, observer bias is often inherent in community science datasets like eBird. We tried to minimize observer bias by filtering data by observer effort, and the inclusion of human modification as a covariate, which accounts for the possibility that checklist observations were proportional to the number of persons residing in the vicinity (i.e., greater sampling effort in highly-modified environments; Humphreys et al., 2019). Lastly, climate change will likely interact with human modification to exacerbate patterns of novelty and homogenization in the future, by selecting for species and populations capable of withstanding more extreme environments (Piano *et al*., 2017; Latimer *et al*., 2018; Williams *et al*., 2020), but the influence of climate change was not accounted for in our analyses.

### Conclusions

We found evidence for homogenization of bird communities in highly human-modified environments at the continental scale in the U.S. but at regional scales, spatial variation in bird community compositions maintained similar levels within both natural and modified habitats. This finding reinforces the growing body of evidence that human-modified landscapes can contribute to broader conservation goals by providing complementary resources for generalist and more synanthropic species capable of persisting in anthropogenic habitats (Kennedy et al., 2017), particularly when impacts can be reduced by less intensive land-use practices (Kremen & Merenlender, 2018; Sol *et al*., 2020). Indeed, cumulative human modification had a significant influence on assemblage turnover within most regions, whereas the proportion of novel assemblages varied widely among regions. By considering assemblage turnover in terms of geography and environmental drivers across different regions, we can unpack some of the mechanisms behind this variation in biotic novelty and its implications. For instance, we found a strong influence of proxies for historical and natural disturbance influences on present-day assemblage turnover in response to human-modification of landscapes, such that the degree of novelty increased along anthropogenic gradients, particularly in regions where past disturbances were low and Euro-American settlement occurred later. Therefore, continued human-modification may result in further widespread human-driven homogenization of bird communities, particularly in the western U.S., which could lead to the loss of valuable ecosystem services and functions.

## Acknowledgements

We are grateful to the thousands of volunteers who contributed observations to eBird, as well as the eBird team at Cornell Lab of Ornithology for curating and maintaining this open-access resource. This work is supported by the National Science Foundation’s Integrated Graduate Education and Training (IGERT) program under award DGE-1144752 and USDA’s National Institute of Food and Agriculture Education and Workforce Development grant 2019-67012-29720.

## Notes

### Competing Interest Statement

The authors have declared no competing interest.

