## Appendix S1 for "Rising novelty and homogenization of breeding bird communities in the U.S."

Figure S1. Sampling coverage for each BCR and human modification type (Natural versus Modified) computed using the improved coverage estimator from Chao and Jost (2012). In all cases, sampling completeness was >95% suggesting near complete inventory of the bird communities within each category and BCR combination.

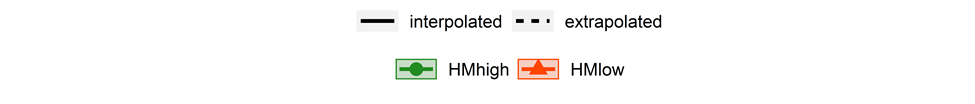

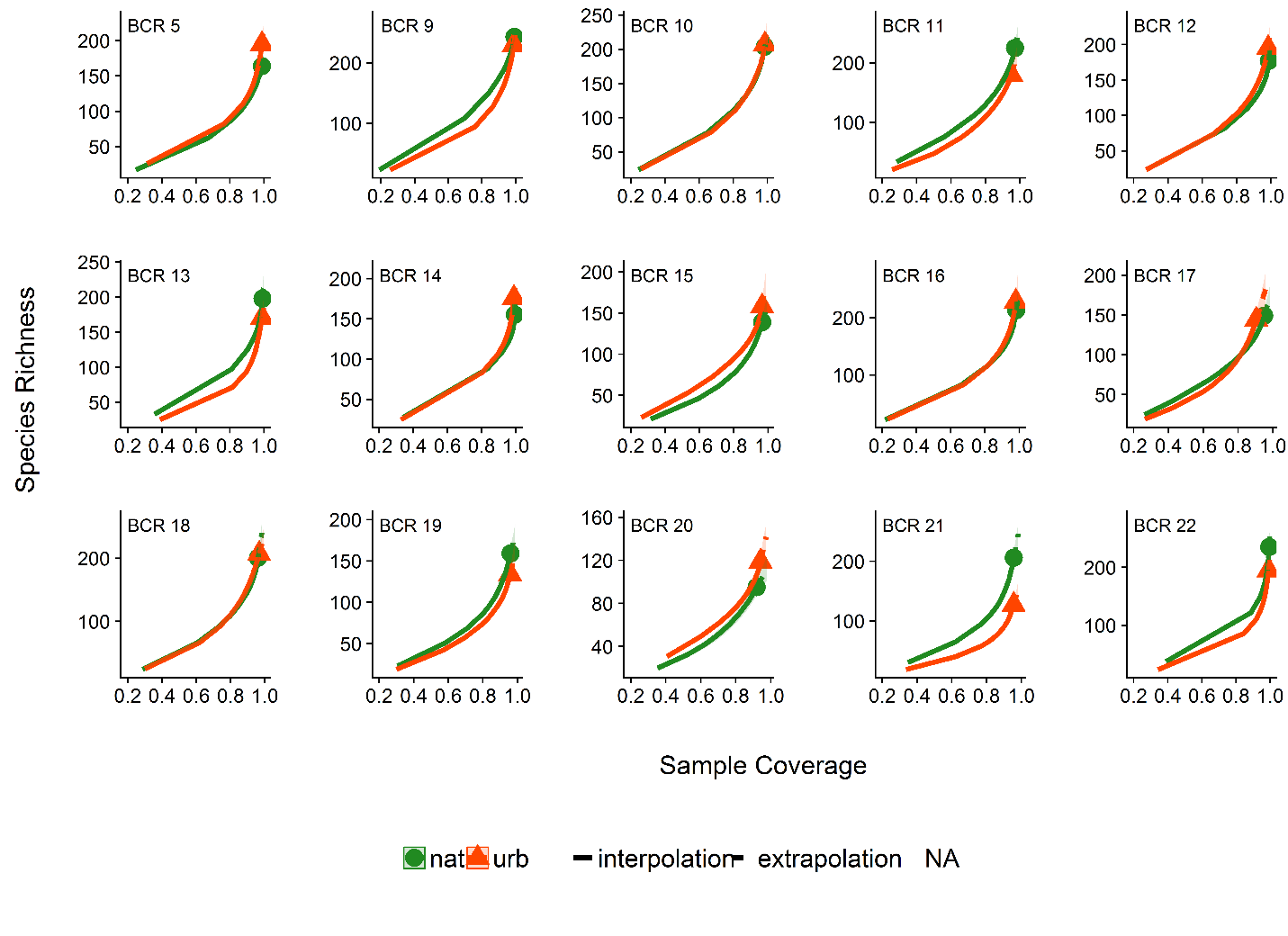

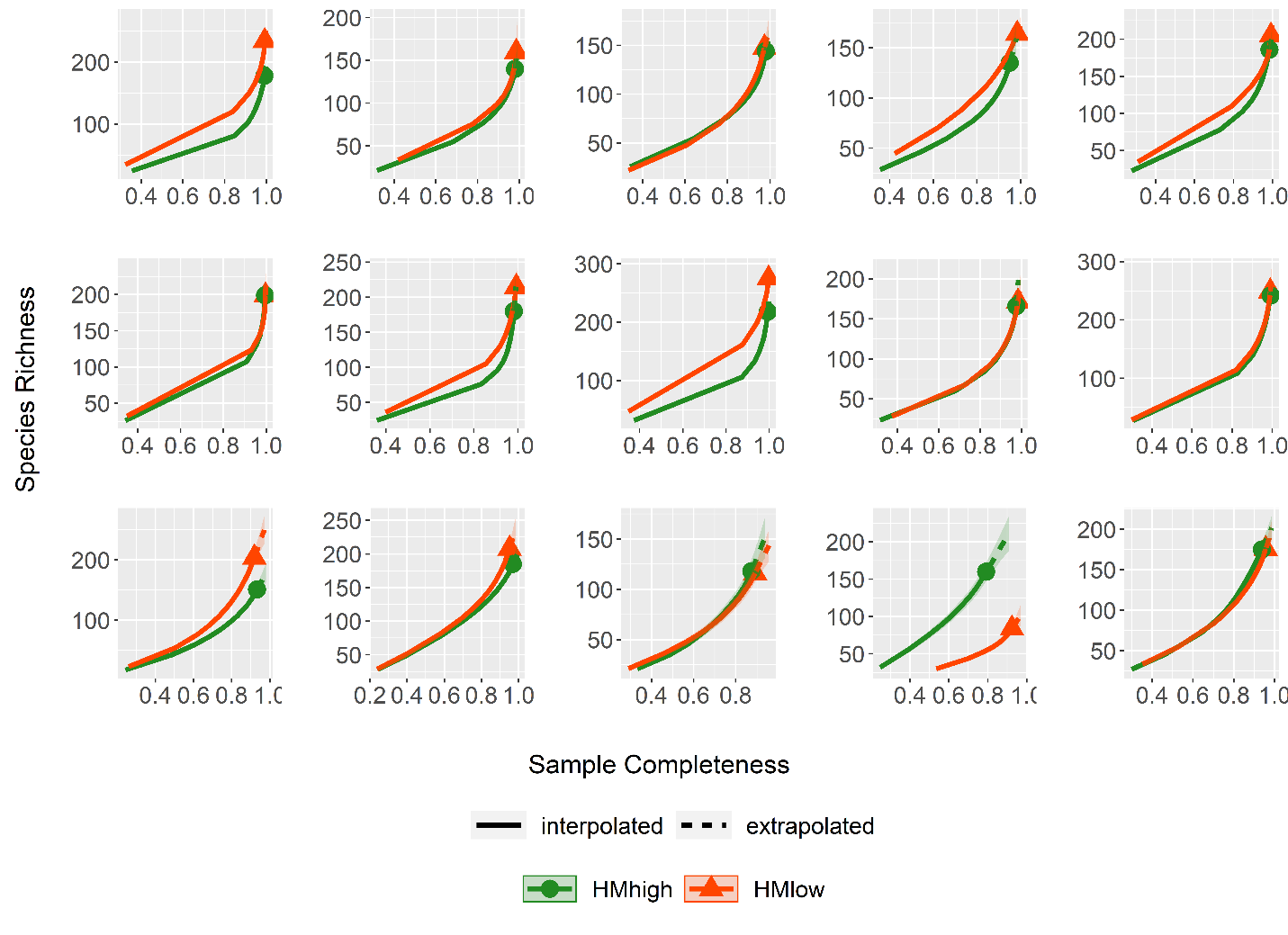
Figure S1 continued

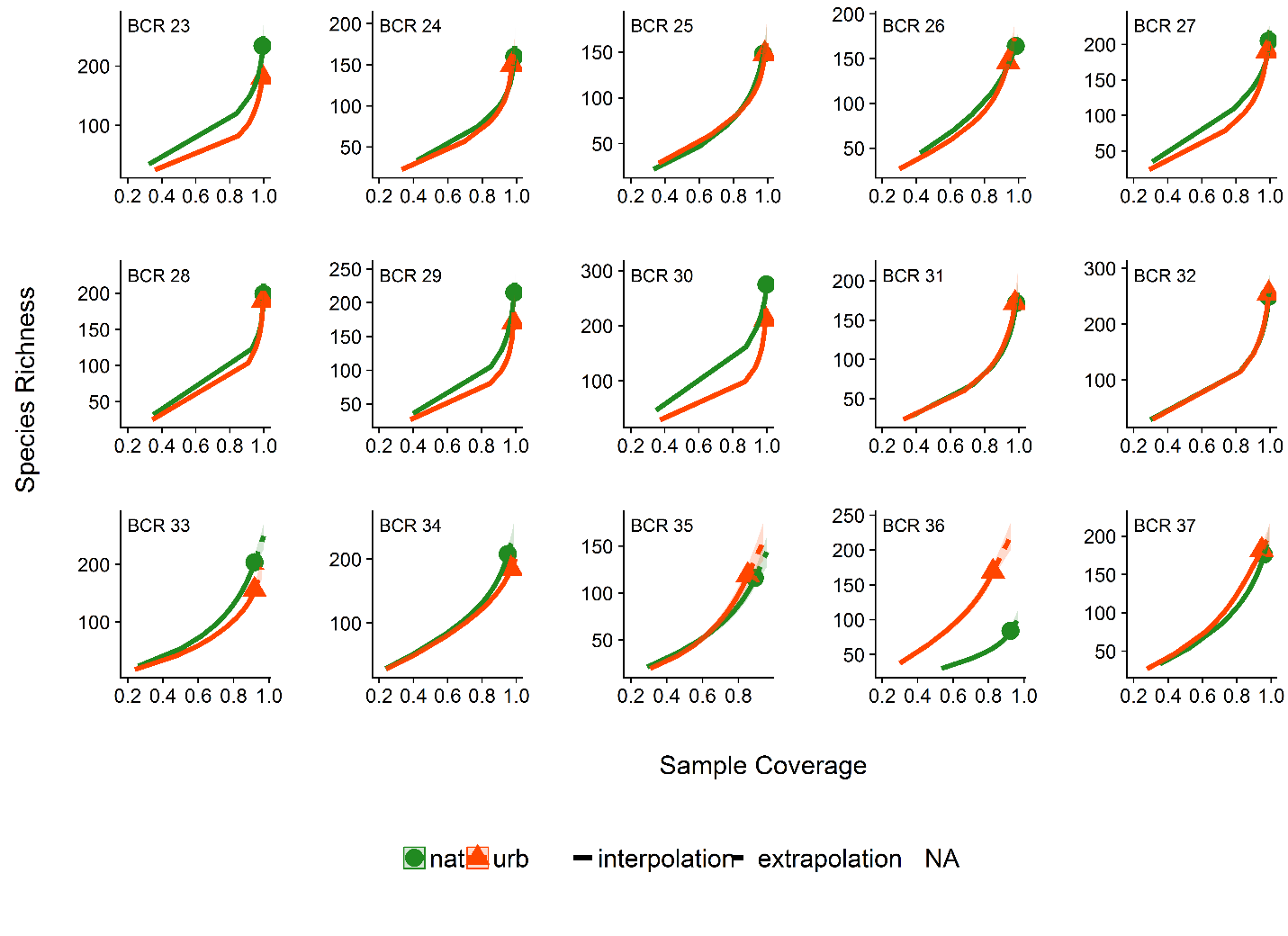

Table S1: Names and numbers of each Bird Conservation Region (BCR); Total number of species pixels (N_sp_pixels) within each BCR; Total number of HM_high_ and HM_low_ pixel assemblages within each BCR (HM_high_ = 95^th^ percentile of human modification values; HM_low_ = 5^th^ percentile of human modification values within each BCR); the proportion of community analogs within each BCR; and the degree of novelty between HM_high_ and HM_low_ assemblage pixels within each BCR. Novelty was calculated as the inverse of proportion of community analogs and re-scaled between 0 (corresponding to low) and 1 (corresponding to high) novelty.

| BCR_Name | BCR | N_sp_pixels | N_pix_HM_high_ | N_pix_HM_low_ | Prop analogs | Novelty |
| --- | --- | --- | --- | --- | --- | --- |
| NORTHERN PACIFIC RAINFOREST | 5 | 5028 | 139 | 139 | 5.9 | 0.67 |
| GREAT BASIN | 9 | 6084 | 38 | 38 | 4.8 | 0.82 |
| NORTHERN ROCKIES | 10 | 5151 | 18 | 18 | 6.5 | 0.59 |
| PRAIRIE POTHOLES | 11 | 2775 | 58 | 58 | 16.6 | 0.19 |
| BOREAL HARDWOOD TRANSITION | 12 | 4332 | 46 | 46 | 7.1 | 0.54 |
| LOWER GREAT LAKES/ ST. LAWRENCE PLAIN | 13 | 4321 | 47 | 47 | 14.7 | 0.22 |
| ATLANTIC NORTHERN FOREST | 14 | 4364 | 55 | 55 | 4.0 | 1.00 |
| SIERRA NEVADA | 15 | 1524 | 90 | 90 | 17.3 | 0.18 |
| SOUTHERN ROCKIES/COLORADO PLATEAU | 16 | 4972 | 94 | 94 | 10.1 | 0.36 |
| BADLANDS AND PRAIRIES | 17 | 1391 | 236 | 236 | 29.0 | 0.08 |
| SHORTGRASS PRAIRIE | 18 | 2799 | 113 | 113 | 10.6 | 0.34 |
| CENTRAL MIXED GRASS PRAIRIE | 19 | 2061 | 37 | 37 | 19.9 | 0.15 |
| EDWARDS PLATEAU | 20 | 680 | 26 | 26 | 60.5 | 0.00 |
| OAKS AND PRAIRIES | 21 | 1921 | 111 | 111 | 17.6 | 0.17 |
| EASTERN TALLGRASS PRAIRIE | 22 | 7103 | 102 | 102 | 14.4 | 0.23 |
| PRAIRIE HARDWOOD TRANSITION | 23 | 8791 | 48 | 48 | 16.0 | 0.20 |
| CENTRAL HARDWOODS | 24 | 3203 | 18 | 18 | 12.6 | 0.27 |
| WEST GULF COASTAL PLAIN/OUACHITAS | 25 | 1263 | 70 | 70 | 17.0 | 0.18 |
| MISSISSIPPI ALLUVIAL VALLEY | 26 | 682 | 29 | 29 | 35.4 | 0.05 |
| SOUTHEASTERN COASTAL PLAIN | 27 | 4732 | 11 | 11 | 9.1 | 0.41 |
| APPALACHIAN MOUNTAINS | 28 | 10166 | 89 | 89 | 5.8 | 0.68 |
| PIEDMONT | 29 | 5195 | 30 | 30 | 7.5 | 0.51 |
| NEW ENGLAND/MID-ATLANTIC COAST | 30 | 5127 | 133 | 133 | 7.7 | 0.49 |
| PENINSULAR FLORIDA | 31 | 2980 | 102 | 102 | 13.2 | 0.26 |
| COASTAL CALIFORNIA | 32 | 4699 | 159 | 159 | 8.6 | 0.43 |
| SONORAN AND MOJAVE DESERTS | 33 | 1674 | 155 | 155 | 15.9 | 0.20 |
| SIERRA MADRE OCCIDENTAL | 34 | 1423 | 149 | 149 | 26.3 | 0.09 |
| CHIHUAHUAN DESERT | 35 | 610 | 45 | 45 | 18.2 | 0.17 |
| TAMAULIPAN BRUSHLANDS | 36 | 440 | 105 | 105 | 9.9 | 0.37 |
| GULF COASTAL PRAIRIE | 37 | 1074 | 66 | 66 | 25.8 | 0.10 |

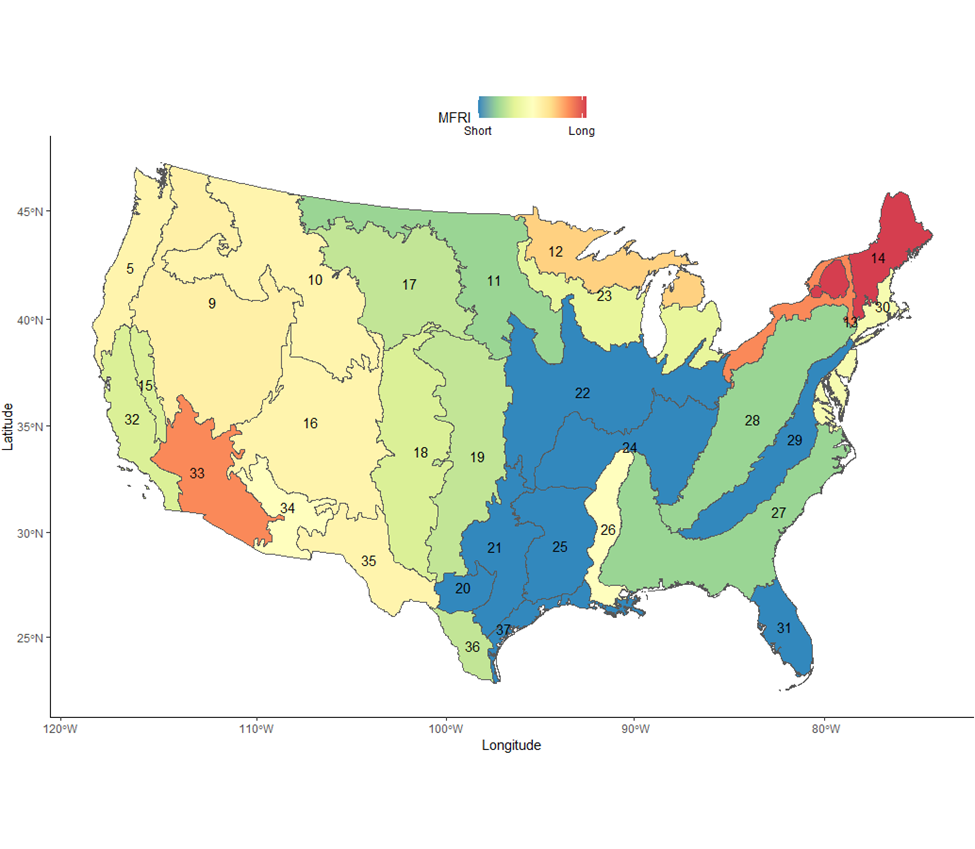

Figure S2. Median value of the Mean Fire Return Interval within each Bird Conservation Region (BCR; gray outline, black number) occurring within the lower 48 U.S. states. Values range from 3 years (blue) to 1000 years (red).

Figure S3. Median year of Native American Land cessions to the U.S. government within each Bird Conservation region (BCR; gray outline, black numbers) throughout the U.S.
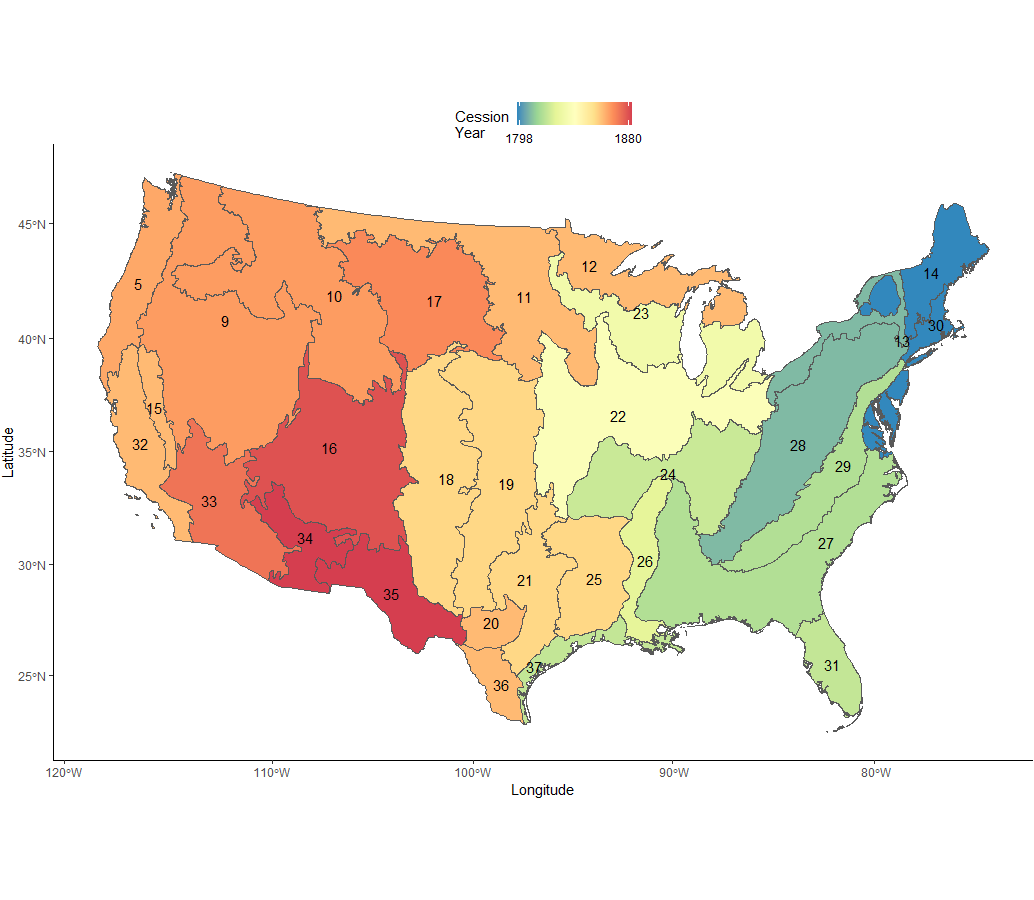

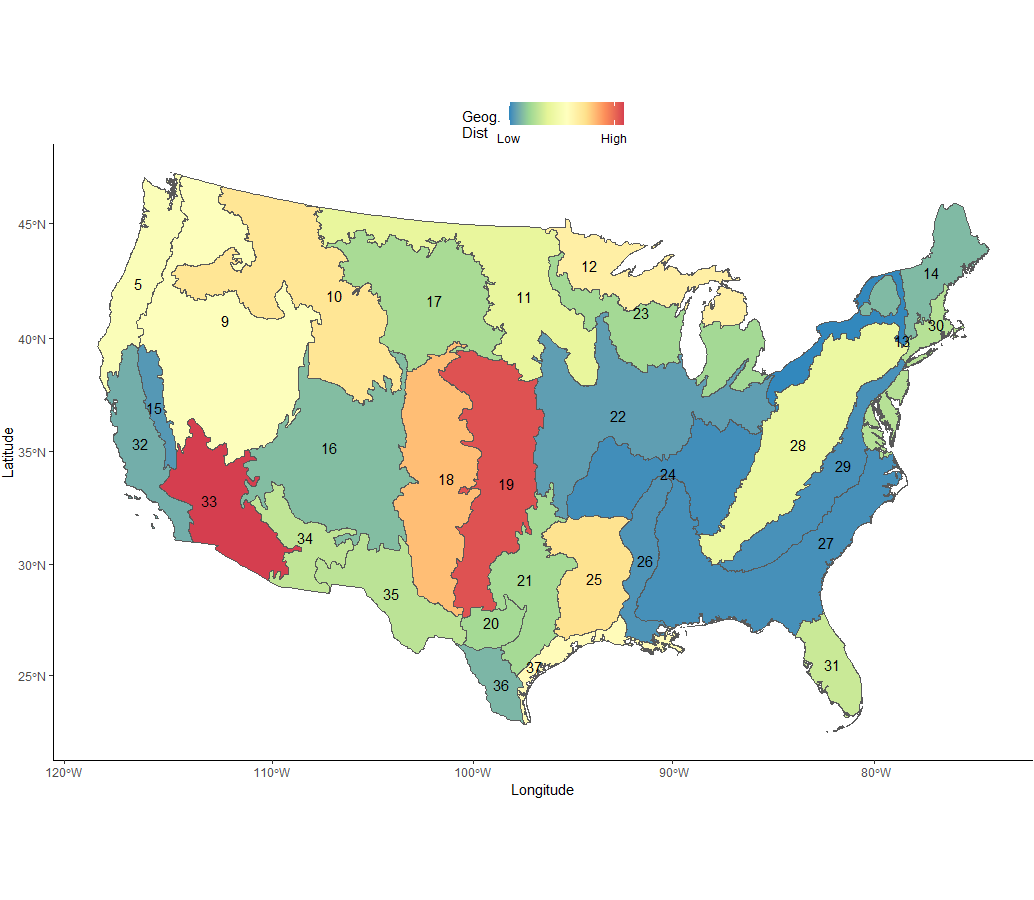

Figure S4. Contribution of geographic distance to avian assemblage turnover mapped to each Bird Conservation Region (BCR; gray outline, black numbering) occurring within the lower 48 U.S. states.
